# Towards a Unified Exact Solution of Rearrangement Small Parsimony for Natural Genomes

**DOI:** 10.64898/2026.06.23.733974

**Authors:** Leonard Bohnenkämper, Daria Frolova

## Abstract

Phylogenetic reconstruction is a fundamental problem in comparative genomics. As a theoretical problem in rearrangement studies, this has been modelled as the Small Parsimony Problem (SPP), in which ancestral genome structures have to be determined minimizing the number of rearrangement events occurring throughout the phylogeny. This problem is of significant interest in microbial and cancer genomics, due to the prevalence and clinical importance of rearrangement events.

Genome structures in this problem are expressed as sequences of *markers*, which are themselves oriented sequence features (such as genes) that abstract from non-structural variations. Recent research has focused on the problem under the natural genomes model, in which arbitrary variations in copy number of markers are allowed. Natural genomes are often studied under the DCJ-indel model, a model which has already been successfully applied to plasmid data. There also exist ILP solutions to a variant of the Small Parsimony Problem under the DCJ-indel model. However, these solutions are limited in their applicability, as they make some critical simplifications for tractability purposes: ancestral marker frequencies and precomputed putative ancestral adjancencies, with their predicted likelihoods, are assumed as input.

This creates multiple problems from both a theoretical and practical perspective. Firstly, this simplification means that not the full state space is searched for a solution, but rather only the subset of genomes with the precomputed putative adjacencies, meaning an optimal solution to the exact SPP is not guaranteed. Secondly, marker frequencies are given externally, without any theoretical guarantees. Thirdly, the method used to precompute adjacencies relies on gene trees, which requires the use of genes as markers, when gene annotation is often unreliable, especially in regions with a lot of rearrangement. Additionally, this restricts the applicability of the approach to sets of genomes that are both divergent and large enough to be able to produce informative gene trees. This is, for example, rarely the case for plasmids, where nucleotide mutations are rarer than rearrangements and genomes are small.

Hence, we revisit the problem to solve the exact SPP by introducing a cost to indel operations, which allows us to compute ranges of marker frequencies and derive theoretical results, that allow us to reduce the solution space that the ILP searches without sacrificing optimality. We show that this makes the problem tractable for the case of small and recently related genomes, first on simulated genomes, and then on a set of pathogenic plasmids which represent a realistic use case for the method.

## 1 Introduction

The problem of parsimonious ancestral reconstruction with a given tree is known as the *Small Parsimony Problem (SPP)*. In general, SPP assumes a given tree with known states at the leaves and a cost function penalising changes between states. It then asks to find ancestral states minimising the total cost in the tree. SPP is solved by Fitch’s method for nucleotide substitutions under unit cost [17]. However, genomic change is not exclusively characterised by substitutions, and structural changes like gene rearrangements and insertions/deletions (indels) of sequence play a significant role in evolution – notable examples include microbial plasmids [23], bacteria [10] and fungi [26], both chromosomal and extrachromosomal DNA in cancer [13, 22], and eukaryotic mitochondrial DNA [2, 31].

To quantify such structural changes various combinatorial rearrangement models have been suggested, in which genomes are treated as sequences or oriented markers, where markers correspond to some sequence features (e.g. genes or synteny blocks). We distinguish here between *singular genomes*, in which each marker occurs at most once and *natural genomes*, for which no restrictions pertaining to copy number is made. In the context of rearrangement models, SPP then asks to reconstruct ancestral marker orders which minimise the total number of rearrangements under a given model.

As a fundamental reconstruction problem, the small parsimony problem has been studied under various rearrangement models. Notably, even for singular genomes, the problem is already NP-hard for only three leaves (median of 3 problem) under complex models, such as the Double-Cut-And-Join model [28] or the Reversal model [8]. In fact, the Problem is only known to be polynomially solvable for the simple Single-Cut-or-Join model [16], which however is not able to handle arbitrary natural genomes.

One model that has been successfully extended to handle natural genomes is the *Double-Cut-And-Join and indel (DCJ-indel) model* [7, 4, 3]. The rearrangement operations in this model are Double-Cut-And-Join (DCJ) – a general rearrangement operation able to mimic reversals, translocations and transpositions – as well as insertions and deletions of one or multiple markers at once. As determining the minimum number of DCJs and indels between a pair of natural genomes is already NP-hard [4], the solutions to this problem [4, 3] are based on an Integer Linear Program (ILP) [25].

The first attempts at a solution to DCJ-indel SPP for natural genomes were undertaken by Doerr and Chauve [12] and Bohnenkämper et al [6]. However, owing to the computational complexity of the problem, these solutions rely on several simplifications. Firstly, the copy number of each marker at ancestral nodes is assumed as input. Secondly, a collection of putative ancestral adjacencies is precomputed, and the ILP searches only within the space of these previously inferred adjacencies, rather than the full space of adjacencies. Thirdly, these precomputed adjacencies are given a weight, and this weight is incorporated into the objective function of the ILP. Although these simplifications help make the problem tractable, it introduces several critical limitations. Since not the full space of adjacencies is searched, it does not truly solve the SPP, and an optimal solution on the full space is not guaranteed. Additionally, the current method used to precompute ancestral adjacencies and their weights relies on gene trees [14]. Gene annotation is often unreliable, especially in regions with structural variation (e.g. even two identical genomes can be annotated differently due to genes being missed), making genes difficult to use as markers in practice [18, 29]. For recently related or very small genomes there is also little SNP information to build gene trees from, making them insufficiently informative for the problem, and therefore limiting its applicability to between-species comparisons.

Naively, one can still achieve a solution to the exact DCJ-indel SPP, and avoid reliance on gene trees, by choosing weights of zero and inputting the entire space of adjacencies. However, this requires knowledge of the correct copy number for each internal node, for which so far no theoretical result exists. Moreover, in practice, such large input data cannot be handled by ILP solvers efficiently, as we demonstrate in Section 4.

The remainder of this manuscript is organised as follows. In Section 2, we give basic definitions and detail previous work. In Section 3.1, we discuss how to parsimoniously infer marker frequencies using a Fitch-like approach. In Sections 3.2 and 3.3, we introduce a way to restrict the solution space, such that at least one optimal solution under the DCJ-indel model is preserved. In Section 3.4, we introduce a way to use strategies like weighting adjacencies as a runtime heuristic as opposed to a correctness heuristic in an ILP context. We evaluate our solution in Section 4 and discuss our findings in Section 5.

## 2 Background

In rearrangement studies, one often abstracts from the concrete sequence content to the higher level of synteny blocks or *markers*. Formally, a marker *m* is a symbol from a (large) alphabet. To represent strandedness, markers are assigned an *orientation*, i.e. +*m* or *− m*. We then speak of an *oriented marker*. A *chromosome* is then modeled as a circular or linear string over the alphabet of oriented markers and a *genome* as a collection of chromosomes.

In our notation, we write a circular string in round and a linear string in square brackets. For example, *E* ={ [+1*−* 2*−* 1], (+3), (+3) }is a genome with a linear and two circular chromosomes. Note that we do not restrict the number of occurrences of a chromosome or marker in our definitions. Also note that inversions of chromosomes are considered equivalent, i.e. [+1 + 2 + 3] = [*−* 3 *−* 2 *−* 1] as well as rotations of circular chromosomes, i.e. (+1 *−* 2) = (*−* 2 + 1).

We define the *universe of natural genomes U* as the set of all possible genomes over a given marker alphabet. Observe that *U* is infinite even though the underlying marker alphabet considered is finite.

The *Small Parsimony Problem* is a reconstruction problem, in which ancestral states are reconstructed for a known tree and known extant genomes in such a manner that the minimum number of evolutionary events is needed to “explain” the resulting phylogeny.

Formally, a binary tree *T* is a connected acyclic graph, such that each node has at most three neighbors. A node is called a *leaf* if it has degree 1, otherwise it is an *internal node*. Sometimes it is helpful to examine only a part of the tree by considering a *subtree* of *T* . To that end, removing some edge (*u, r*) ∈ *E*(*T*) splits the tree into two components. Formally, the *subtree* with *root r* is then the component containing *r*. In the Small Parsimony Problem, each leaf represents an extant genome whereas each internal node represents the most recent common ancestor of all leaves in one of the subtrees of which it is the root.

Given a rearrangement distance measure *d*, the Small Parsimony Problem in the context of natural genomes can then be simply defined as follows.

### Problem 2.1

(Small Parsimony Problem for Natural Genomes). *Given a binary tree T with edges E*(*T*) *and genome labels L*_*u*_ *∈ U for each leaf u, find labels L*_*v*_ *∈ U for each internal node v, such that*

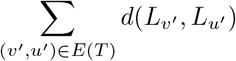

*is* minimised *under distance d*.

### 2.1 Previous Work

In this subsection, we summarise previous work, which is necessary to derive the results of our contribution. We begin by discussing the DCJ-indel distance in Section 2.1.1 and the previous approach by Doerr and Chauve to solve a restricted version of SPP in Section 2.1.2.

#### 2.1.1 DCJ-indel distance for natural genomes

Here we summarise previous results on an important part of the distance model *d* that we will later use in Problem 2.1. Rearrangement distances between two genomes *A, B* ∈ *U* are often defined as the minimum number of rearrangement operations under a chosen model needed to transform *A* into *B*.

The problem of determining this minimum number of rearrangements between two genomes is known as the *distance problem*. To that end, it is often useful to disambiguate between different occurrences of the same marker. We thus presume that there is some arbitrary labelling *A*^*l*^ for each genome *A* that adds a unique sub-indentifier *i* to each occurrence of a marker *m*, obtaining *m*_*i*_. We call *m*_*i*_ a *marker occurrence* and *A*^*l*^ a *labelled genome*. For example, *E*^*l*^ = {[+1_1_ *−* 2_1_*−* 1_2_], (+3_1_), (+3_2_) }would be one of multiple labellings for the genome *E* introduced previously. From now on, we presume each genome is a labelled genome.

Each (labelled) genome can be equivalently described by its *adjacencies*. For each marker occur-rence *m*_*i*_ in *A*, we denote its beginning by 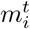(tail) and its end by 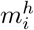(head), i.e. 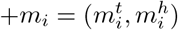 and 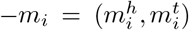. We call 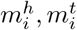 *extremities*. An *adjacency* is then a pair of extremities 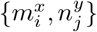, *x, y ∈ {t, h}*, such that the corresponding marker occurrences *m*_*i*_ and *n*_*j*_ are neighbor-ing on a chromosome and 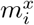and 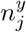are adjacent. For example, in *E*^*l*^ we have the adjacencies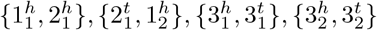. In linear chromosomes, the first extremity of the first marker and second extremity of the last marker are not adjacent to any other extremity. For mathematical convenience we thus define a *telomeric adjacency* 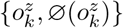 for such an extremity *o*^*z*^ for a unique artificial extremity 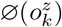. 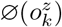 is called a *pseudo-cap*. For example, in *E*^*l*^ we have the telomeric adjacencies 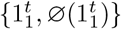 and 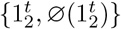. When the concrete extremities are irrelevant, we use the shorthand notation 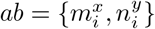 to denote the adjacency as an arbitrary pair of extremities *a* and *b*. Note that the order of *a* and *b* is still arbitrary, i.e. *ab* = *ba*.

A relatively simple but powerful rearrangement model is the DCJ-indel model [7]. It was first introduced on *singular genomes*, a finite subset *S⊂ U* of all natural genomes. In a singular genome each marker may occur at most once. To this end, we define the *copy number ϕ*(*A, m*) of marker *m* as the number of times marker *m* occurs in genome *A* as some occurrence *m*_*i*_. For example, earlier in *E*^*l*^ we have *ϕ*(*E*^*l*^, 1) = *ϕ*(*E*^*l*^, 3) = 2, *ϕ*(*E*^*l*^, 2) = 1. We can then define the set of singular genomes as *S* := {*A ∈ U* |∀ *mϕ*(*A, m*) ≤ 1} . Moreover, we say a marker *m* with *ϕ*(*A, m*) = 1 and its extremities are *singular* in *A*.

Within the DCJ-indel model, the following operations are permitted:

- A *Double-Cut-And-Join (DCJ)* transforms the adjacencies of a genome in one of the following ways:

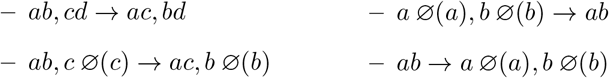
- An *insertion or deletion* (indel), in which a segment of one or multiple markers is inserted or deleted at any position in the genome.

In order to prevent “free lunch” scenarios, in which chromosomes are simply deleted as a whole, the original formulation in [7] allows only to insert or delete non-shared markers between the two genomes. The DCJ-indel distance 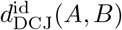 for two singular genomes *A, B ∈ S* is then the minimum number of DCJ and indel operations needed to transform one genome into the other.

This distance can be calculated in linear time using the *relational diagram*. Given two singular genomes *A* and *B*, the *Relational Diagram (RD) R*(*A, B*) is a graph (*V, E*). Its set of vertices is *V* := *V* (*A*) ∪ *V* (*B*), where *V* (*A*) has a vertex for each extremity of each marker occurrence of genome *A*, and *V* (*B*) has a vertex for each extremity of each marker occurrence of genome *B*. For any adjacency *ab* in *A*, an *adjacency edge* is added between vertices *a* and *b*, and similarly for any adjacency in *B*. Additionally, for every shared marker *m*, any 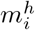in *V* (*A*) is connected to any 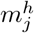 in *V* (*B*) by an *extremity edge*, and similarly any 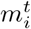 in *V* (*A*) to any 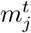 in *V* (*B*). An example for a RD is given in Figure 1.

**Figure 1.**
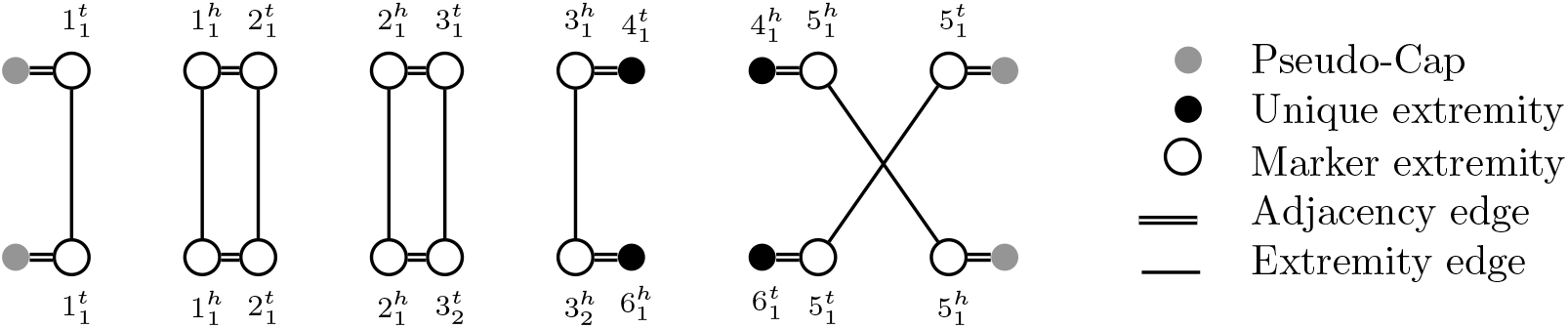
Relational Diagram (RD) for singular genomes *A* = *{*[+1_1_ + 2_1_ + 3_1_ + 4_1_ *−* 5_1_]*}* and *B* = *{*[+1_1_ + 2_1_ + 3_1_ *−* 6_1_ + 5_1_]*}*.

Notably, since each marker occurs at most once in each genome, the RD consists only of simple cycles and paths. Paths can either end in a pseudo-cap, in which case we denote the path end by the genome name to which the pseudo-cap belongs in capital letters (*A* or *B*) or in an extremity unique to one of the genomes, in which case we denote the path end by the genome name in lowercase letters (*a* or *b*). All combinations of these path ends yield 10 different path types, the counts of which in a given Relational Diagram we denote accordingly by *p*_*AA*_, *p*_*AB*_, *p*_*BB*_, *p*_*Aa*_, *p*_*Ab*_, *p*_*Ba*_, *p*_*Bb*_, *p*_*aa*_, *p*_*ab*_, *p*_*bb*_. We write the number of cycles as *c*. For example, in Figure 1, we have *c* = 2, *p*_*AB*_ = 1, *p*_*ab*_ = 1, *p*_*Ab*_ = 1, *p*_*Ba*_ = 1 with all other counts being 0.

*Circular singletons* are structures that are not represented in the graph, but still necessary to calculate the DCJ-indel distance. A circular singleton is a circular chromosome consisting only of markers unique to one genome. We denote the number of circular singletons by *s*. In [3], the following result is derived.

##### Theorem 2.2

(adapted from [3]). *For two singular genomes A, B, the DCJ-indel distance is*

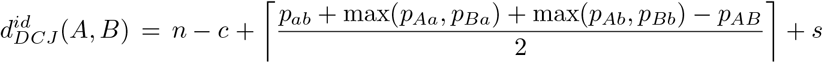

*with n the number of* shared markers.

The distance problem of natural genomes can be reduced to the distance problem of singular genomes by employing a *matching*, which informs for a marker *m* which occurrence *m*_*i*_ in *A* should be considered the same as *m*_*j*_ in *B*. A matching between genomes *A* and *B* is defined as a set of pairs *M* ={ (*m*_*i*_, *m*_*j*_) |*m* shared marker, *m*_*i*_ occurrence in *A, m*_*j*_ occurrence in *B*}, such that each marker occurrence appears at most once in *M* .

Given a matching for each marker in *A* and *B*, we can compare them as singular genomes. For example, given (*m*_*i*_, *m*_*j*_) *∈ M* can then rewrite both *m*_*i*_ in *A* and *m*_*j*_ in *B* to the same unique character *u*^*ij*^, and rewrite any occurrence not in *M* to a character that does not occur in the other genome. We call the resulting genomes *A*^*M*^, *B*^*M*^ the *singularisations* of *A* and *B*. The distance between two natural genomes can then be defined as the minimum distance between any two singularisations obtained by a matching.

However, not all matchings are meaningful. For example, choosing the empty matching would lead to all chromosomes being inserted or deleted, which under the DCJ-indel distance may actually be optimal, since it requires only one (indel) operation per chromosome. One way to resolve this issue is to require the matching to be *maximal*, i.e. such that it cannot be extended. In this case, this is equivalent to the matching having maximum cardinality. Formally, we write M(*A, B*) as the set of all possible matchings between two genomes *A, B* and 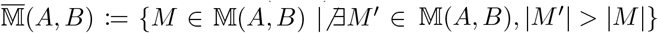 as the set of *maximal matchings*.

In [4, 12, 6], the distance measure 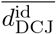for two natural genomes *A, B* is defined as the minimum DCJ-indel distance between any singularisations obtained by a maximal matching:

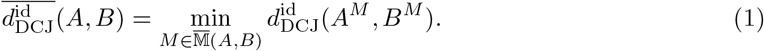

We call this distance model the *restricted* DCJ-indel distance.

#### 2.1.2 Linearising degenerate genomes

The first attempt to solve the small parsimony problem was undertaken by Doerr and Chauve in 2021 [12]. In their formulation, the authors restrict Problem 2.1 in two critical ways: (I) It is assumed that the copy number *ϕ*(*L*_*v*_, *m*) of the genome label *L*_*v*_ *∈ U* internal node *v* is given for each marker *m*. This restricts the infinite universe *U* to a finite subset of candidate genomes. (II) It is assumed that even within that restricted subset, only some adjacencies are permitted, further restricting the space of possibilities.

The first assumption about marker copy numbers was relaxed in [6], allowing for a range of given copy numbers *l*(*v, m*) ≤ *ϕ*(*L*_*v*_, *m*) ≤*h*(*v, m*) for the genome label *L*_*v*_ at internal node *v*. Nonetheless, this solution still relies on external heuristics to derive copy number counts and candidate adjacencies, which inform the finite candidate set that is explored. The advantage of this formulation is that the solution space with given input copy numbers and adjacencies can be encoded as a *degenerate genome*.

##### Definition 2.3

(adapted from [12]). *A* degenerate genome *is a set of adjacencies D that satisfies the following conditions:* 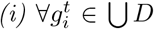, *there exists also extremity* 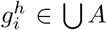 *and vice versa, and (ii) each telomeric extremity* 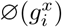 *is used only in the adjacency* 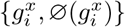.

The solution space spanned by a degenerate genome *D* is then each genome that has only adjacencies from *D*. Such a genome is called a *linearisation of D*. Formally, a *linearisation* of degenerate genome *D* is a genome *A*, such that each adjacency 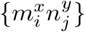 of *A* is part of *D*, i.e.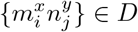.

In the original formulation [12], each extremity in *D* must be used in the linearisation *A*. However, since the ILP solution in [6] allows for variant copy numbers, we choose a slight reformulation that explicitly encodes the upper and lower bounds for each marker. The variant of SPP solved in [6] can then be formulated as follows.

##### Problem 2.4

(adapted from [6]). *(Weighted Small Parsimony Linearisation Problem) Given a binary tree T with edges E*(*T*), *genome labels L*_*u*_ *∈ U for each leaf u, degenerate genomes D*_*v*_ *for each internal node v, a weighting function w*_*v*_ *for adjacencies of each degenerate genome D*_*v*_, *lower and higher bounds l*(*v, m*), *h*(*v, m*) *for each marker m at each internal node v and a parameter α ∈* [0, 1], *find a linearisation L*_*v*_ *for each degenerate genome D*_*v*_, *such that l*(*v, m*) ≤ *ϕ*(*L*_*v*_, *m*) ≤ *h*(*v, m*), *minimising*

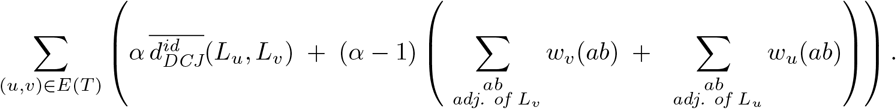

Observe that what is optimised in this problem formulation is not the restricted DCJ-indel distance, but, depending on *α*, also a weight bias *w*_*v*_ toward the input adjacencies at each node *v*. These weights are part of the input and result from confidence scores of the adjacency prediction for ancestral genomes. The consequence of this formulation is that the set of degenerate genomes and weights associated with each of their adjacencies are assumed to be given, and must be precomputed before solving Problem 2.4. Since these are derived externally and without any theoretical guarantees, this formulation does not solve the SPP for natural genomes as formulated in Problem 2.1.

Note however, that if one could guarantee that the optimal solution is encoded in the degenerate genomes *D*_*v*_ and set *α* = 1, Problem 2.4 could be used to solve to Problem 2.1. In the following, we will develop the theoretical background that allows us to do exactly this. We will show how to restrict the copy number of each marker to guarantee all optimal solutions are still represented and derive results that help to restrict the space of possible adjacencies while guaranteeing at least one optimal solution can be found.

### 2.2 Problem Definition

Besides restricting the infinite solution space *U* to the finite solution space expressible by degenerate genomes, the reason why marker copy numbers *ϕ* are assumed to be given in [12] and [4], is that using 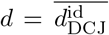 in Problem 2.1 would inherently cause problems due to some adverse properties of the restricted DCJ-indel distance. Importantly, this distance is not a metric, because it violates the triangle inequality [7]. The most striking example for this is choosing any two unichromosomal genomes 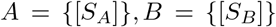 for any oriented strings *S*_*A*_ and *S*_*B*_ over the marker alphabet that have distance 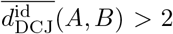. If we now regard the empty genome *C* = *{}*, we observe that 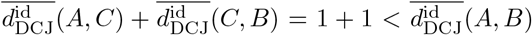 as only a single segmental deletion or insertion is necessary to transform *A* into *C* or *C* into *B*. The empty genome is thus almost always among the genomes with the lowest total restricted DCJ-indel distance to any set of genomes. Thus, solving Problem 2.1 simply setting 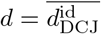 will result with high likelihood in empty or near-empty ancestral genomes.

However, instead of restricting marker multiplicities externally, we opt to instead restore the triangle inequality for the DCJ-indel distance. To that end, we adapt the solution suggested by Braga, Willing and Stoye in the original publication of the DCJ-indel model [7]. There the authors propose to use a surcharge *δ* for each inserted or deleted marker between two singular genomes. Essentially, an indel, instead of having fixed cost 1, instead has cost 1+*δl* where *l* is the length of the inserted or deleted segment. For singular genomes, this cost can then be calculated after determining the default DCJ-indel distance by simply examining the non-shared markers. Defining the non-shared markers as *A△B* := {*m* | *ϕ*(*A, m*) = 1, *ϕ*(*B, m*) = 0} ∪ {*m* | *ϕ*(*A, m*) = 0, *ϕ*(*B, m*) = 1}, this distance is then 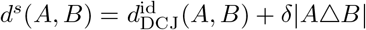.

Note that this result is restricted to singular genomes *S*. In order to apply it to all natural genomes *U*, we can generalise as follows.

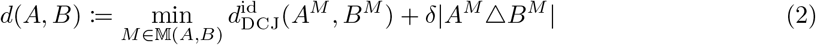

Importantly, we allow any matching *M ∈ M*(*A, B*) here, not just maximal matchings. Observe that again *d* is equivalent to the minimum cost of transforming *A* into *B* when each DCJ has cost 1 and each indel of length *l* costs 1 + *δl*, but with no restrictions on which markers can be inserted or deleted – even shared markers can be part of indel operations. We then arrive at the following.

#### Theorem 2.5

*d is a metric on U for any δ ∈* ℝ, *δ ≥* 0.

*Proof* Identity and symmetry follow directly from the identity and symmetry of the DCJ-indel distance and identity and symmetry of |*A*^*M*^△ *B*^*M*^| . What remains to be shown is the triangle inequality, which follows directly from *d* being a transformation distance. Let thus *A, B, C*∈ *U* . Observe that if transforming *A* into *C* has cost *d*(*A, C*) and *C* into *B* has cost *d*(*C, B*), then there is a way to transform *A* into *B* with cost *d*(*A, C*) + *d*(*C, B*). Thus, *d*(*A, C*) + *d*(*C, B*) *≥ d*(*A, B*).

In principle, we can thus choose any *δ* for our formulation, but choosing a small *δ* will lead to the same problems as the pure DCJ-indel distance. In previous work on natural genomes, these problems are circumvented by allowing only maximal matchings [4, 12, 6].

Note maximising the matching size |*M*| is equivalent to minimising |*A*^*M*^ *△B*^*M*^ |. In the context of our distance model *d*, this is then equivalent to regarding *δ → ∞*, so that any difference in *δ*|*A*^*M*^ *△B*^*M*^ | is greater than any difference in 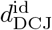, enforcing as many markers as possible are matched. This means for our formulation we can first minimise *δ*|*A*^*M*^ *△B*^*M*^ | and then minimise 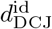 under the maximal matching model.

## 3 Methods

Our method rests on reducing Problem 2.1 to Problem 2.4, that is, we find degenerate genomes that guarantee to encode the optimal linearisations from the universe *U*. To that end, knowing the copy number for each marker is essential. Since we presume *δ*| *A*^*M*^△ *B*^*M*^| is the dominant term in *d* and can therefore be optimised in advance, we first derive marker copy numbers along the tree based on this cost in Section 3.1. Once we know the copy number bounds *l*(*v, m*), *h*(*v, m*) for each marker *m* at each internal node *v*, we can then create the degenerate genome *D*_*v*_ consisting of all possible adjacencies for the extremities in 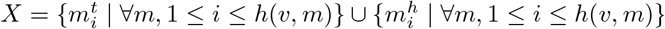, i.e. *D*_*v*_ = {{ *x, y*}| *x* ∈ *X, y* ∈ *X } ∪* {(*x*, ∅(*x*)) | *x ∈ X}* . We call such a genome *fully degenerate*.

In principle, this then allows us to solve SPP by solving a variant of Problem 2.4. We do this by. setting *α* = 1 and enforcing in the ILP of [6] that the total cost contributed by 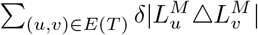 in the tree is exactly the minimum calculated beforehand.

However, in practice, it makes sense to further restrict the space of possible adjacencies. To accomplish this in a manner that guarantees at least one optimal solution remains, further theoretical results are necessary. We derive these results in Section 3.2.

Moreover, when we consider the full space of adjacencies, this opens up a way to restrict the space of possible matchings, a technique that could not be used in previous solutions to the problem. We describe the relevant theoretical results in Section 3.3.

Finally, we develop in Section 3.4 how we can use aspects of correctness heuristics like the weights used in Problem 2.4 as part of our runtime heuristic to obtain good solutions early in the solving process.

### 3.1 Finding Optimal Marker Frequencies

Since under the our distance function a cost of *δ* is needed for each marker that is not matched to another, changes in copy number of marker *m* between two linearisations *L*_*u*_, *L*_*v*_ that are adjacent in the phylogeny ((*u, v*) ∈ *T*) imply a cost of *δ* |*ϕ*(*L*_*u*_, *m*) *− ϕ*(*L*_*v*_, *m*) | . We can thus infer marker copy numbers under this model, minimising this total cost across the phylogeny. Formally, this is equivalent to a character-state small parsimony problem, where each marker is a character and states are copy numbers, drawn from the alphabet ℕ_0_ and the cost function is *c*(*x, y*) = *δ*| *x− y*| . Since any cost in this model is scaled by *δ*, we can instead equivalently consider the cost function *c*^*′*^(*x, y*) = |*x −y*| .

This is then a Wagner Parsimony Problem, for which ancestral copy numbers are easily computed using the Fitch-like Farris optimisation [15]. An efficient, polynomial algorithm to construct all possible character state ranges is given by Swofford and Maddison in [27].

### 3.2 Optimally Filtering Adjacencies

With the maximum copy number per marker known, we can now build a fully degenerate genome for each internal node that is guaranteed to have all optimal genome labels from *U* for that node as linearisations. However, this solution space is much larger than needed for many problem instances. Particularly, since we are interested in phylogenies with a recent history, distances in subtrees close to the leaves are expected to be small. The following lemma allows us to use this fact to restrict the solution space for adjacencies in these cases. To that end, notice that the adjacencies of a genome *A* are themselves a degenerate genome that permits exactly one linearisation, namely *A*. Therefore, any Lemma that applies to degenerate genomes also applies to genomes in general.

#### Lemma 3.1

*Given three (degenerate) genomes D*_*v*_, *D*_*u*_, *D*_*w*_, *at distinct nodes u, v, w, such that* (*u, w*), (*v, w*) *∈ E*(*T*) *and the maximum distance between any linearisations of D*_*v*_ *and D*_*u*_ *is* 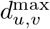 *and D*_*u*_ *⊂ D*_*w*_. *For the optimal linearisations* 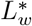 *of D*_*w*_ *and* 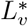 *of D*_*v*_ *holds* 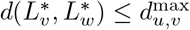.

*Proof* For the sake of contradiction assume 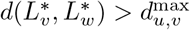. Let (*x, w*) *∈ E*(*T*) with *x≠u, x ≠v*, i.e. *x* is the third node adjacent to *w* in the tree with degenerate genome *D*_*x*_ and optimal linearisation 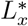. The total cost contributed to the tree at node *w* by 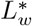 is then 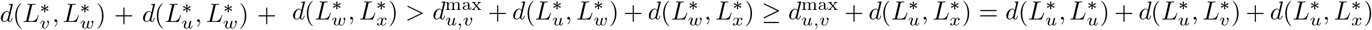. Therefore, replacing 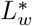 with 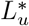as a linearisation of *D*_*w*_ would yield a smaller total cost. 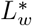 would thus not be optimal, a contradiction.

Most importantly, we can use this result for two adjacent leaves (“cherries”) that have a distance of at most 1.

#### Corollary 3.2

*For two leaves u, v incident at internal node w*, (*u, w*), (*v, w*) *∈ E*(*T*), *such that d*(*L*_*u*_, *L*_*v*_) *≤* 1. *Then the optimal linearisation* 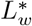*of D*_*w*_ *is L*_*u*_ *or L*_*v*_.

In fact, we can iterate this argument: Since 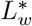 is then either *L*_*u*_ or *L*_*v*_, we can restrict the degenerate genome *D*_*w*_ to only permit *L*_*u*_ or *L*_*v*_ as linearisations. If we then consider two other distinct nodes *x, w*^*′*^ with (*x, w*), (*x, w*^*′*^) *∈ E*(*T*), such that all linearisations of *D*_*w*_, 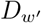have distance of at most one, we can restrict the candidate set *D*_*x*_ as well. As long as the distance between genomes is at most 1, we can remove entire subtrees from the problem without sacrificing optimality.

#### Corollary 3.3

*The optimal linearisation* 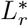 *of D*_*r*_ *of any subtree E*^*′*^ *with root r and leaves L*^*′*^, *such that for all u, v ∈ L*^*′*^ *d*(*L*_*v*_, *L*_*u*_) *≤* 1 *is a genome from {L*_*v*_ | *v ∈ L}*.

We can then remove all nodes of *T* ^*′*^ except *r* from the tree. Moreover, the optimal linearisation for all other nodes can be determined using the Fitch algorithm: In a bottom up pass, linearisation candidates at each internal node are determined and after the optimal linearisation 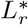 is found, a top down pass decides which linearisation candidate is used.

These results help to keep the solution space small when distances are small. However, even when distances are larger, individual adjacencies often stay preserved within a subtree. The next observation allows us to exploit this fact.

#### Lemma 3.4

*Given (degenerate) genomes D*_*u*_, *D*_*v*_, *D*_*w*_, *D*_*x*_ *for distinct nodes u, v, w, x, such that* (*u, w*), (*v, w*), (*w, x*) *∈ E, and such that any optimal linearisations L*_*u*_, *L*_*v*_ *of D*_*u*_ *and D*_*v*_ *both contain adjacency ab where a, b are singular extremities and D*_*w*_ *is fully degenerate, but any optimal linearisation contains a, b as singular extremities. Then for any optimal linearisation* 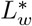 *of D*_*w*_, *there is a linearisation* 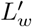 *that contains ab and is co-optimal*.

*Proof* Regard optimal linearisations 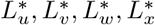 of *D*_*u*_, *D*_*w*_, *D*_*w*_, *D*_*x*_. Assume 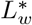 does not con-tain *ab*, but instead *aa*^*′*^ and *bb*^*′*^ with *a*^*′*^ *b, b*^*′*^ *a*. Regard 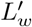, which differs from 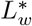 by in-stead containing adjacencies *ab* and *a*^*′*^*b*^*′*^, all other adjacencies being identical to 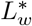 . Observe that 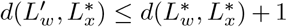, since 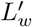 differs from 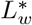 by a single DCJ. Moreover, the components in the Relational diagrams between 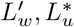 and 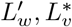 differ from those of 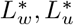 and 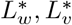 by only an extra cycle (see Figure 2). This means the term *c* (see Theorem 2.2) is increased by one. Since no other component types are changed, all terms based on the graph remain the same. The number of common markers *n* remains also the same. The only term that could have changed otherwise is *s*, the number of circular singletons. However observe that if *a*^*′*^ and *b*^*′*^ are part of a circular singletons, it means their markers incur a cost of *δ* upon being deleted when transforming 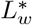 into 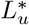 or 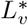 . Since *δ*| *A*^*M*^ △*B*^*M*^ |is the dominant term in *d*, 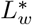 could not have been optimal if they are circular singletons with both 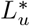 and 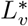 . Let thus w.l.o.g. *a*^*′*^*b*^*′*^ not be a circular singleton for 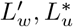. Then, 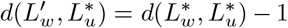 and 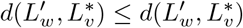. With 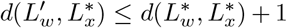, 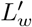 is thus at least co-optimal to 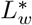 .

Again, this restriction can be propagated through the tree. As long as an adjacency consists of singular extremities and is preserved in the subtree, no alternatives have to be considered. Once duplicates are involved, the solution space is better constrained on the matchings than the adjacencies as we delineate in the next subsection.

**Figure 2.**
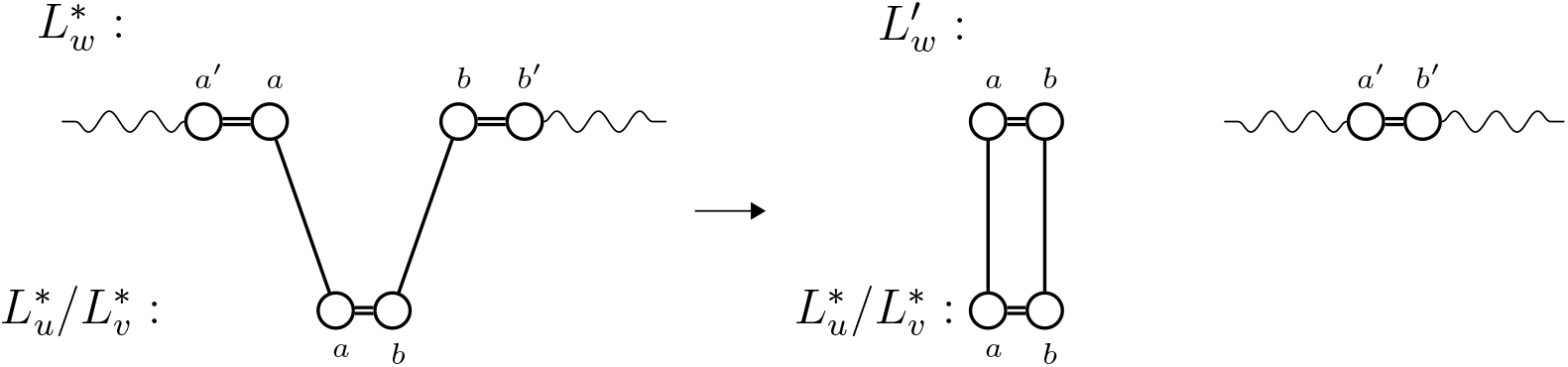
Difference for the Relational Diagrams of 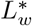 (top left) and 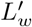 (top right) with 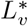 or 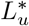 (both bottom). Squiggled lines represent arbitrary simple paths. 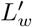 has an extra cycle and otherwise the same graph component types. Particularly, the type of component that *a*^*′*^, *b*^*′*^ are in is unchanged.

### 3.3 Optimally restricting the Matching Space

Note that with duplicates, we are not able to meaningfully restrict the adjacency space unless distances are small (Corollary 3.3). However, because we are using the full adjacency space, we are able to restrict the number of different matchings. To see how this is possible observe the following pair of labeled genomes: *E*_2_ ={ [+1_1_ *−*2_1_ + 2_2_] }, *E*_3_ = {[+1_1_ *−*2_2_ + 2_1_]} . Both are distinct linearisations of the same fully degenerate genome. However, both linearisations represent the same structure – the distance to any other genome *D* is the same, *d*(*E*_2_, *D*) = *d*(*E*_3_, *D*). Only the matching to achieve this distance will be different, i.e. if the matching between *E*_2_ and *D* contains (2_1_, 2_*i*_), the corresponding matching for *E*_3_ and *D* contains (2_2_, 2_*i*_). We can exploit this redundancy to restrict the matching space as follows.

#### Lemma 3.5

*Let D*_*u*_ *and D*_*v*_ *be degenerate genomes. Let marker m occur k times in D*_*u*_ *and n ≥ k times in D*_*v*_, *such that each occurrence in D*_*v*_ *has the same adjacencies, i*.*e*.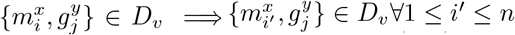. *Then for any linearisations L*_*u*_, *L*_*v*_ *with optimal matching M, there are linearisations* 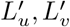, *such that their optimal matching M*^*′*^ *contains only pairs of the form* (*m*_*i*_, *m*_*i*_) *and* 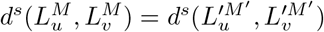.

*Proof* By induction over *x*(*M*) := |*{*(*m*_*i*_, *m*_*j*_) *∈ M* | *i j}*|.

*Induction start* Let *x*(*M*) = 0. Then *M* is already a matching that conforms to the lemma, i.e. 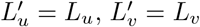 and *M*^*′*^ = *M* .

*Induction step* Let *x*(*M*) *>* 0. Let (*m*_*i*_, *m*_*j*_) *∈ M* with *i≠ j*. Then switch positions of *m*_*i*_ and *m*_*j*_ in *L*_*v*_ to obtain 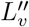. Then replace (*m*_*i*_, *m*_*j*_) in *M* by (*m*_*i*_, *m*_*i*_) and if (*m*_*k*_, *m*_*i*_) *∈ M*, replace it by (*m*_*k*_, *m*_*j*_), obtaining matching *M*^*′′*^. The distance is then the same, 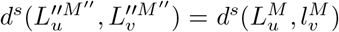, and *x*(*M*^*′′*^) *≤ x*(*M*) *−* 1.

For any internal node *u* with degenerate genome *D*_*u*_ and *n* copies of marker *m*, we can thus find a neighbour (*u, v*) ∈ *E*(*T*) that has *k*≤ *n* copies of *m* and allow only matchings of the type (*m*_*i*_, *m*_*i*_) in the ILP solution presented in [6]. Note that we can do this only once per marker per internal node, that is restricting the matching space for more than one neighbour would lose optimality. For maximum effect, we chose the neighbour with the highest number of matching candidates.

### 3.4 Solving Strategy

While the weighting for Problem 2.4 modulates between an optimisation based on weights and an optimisation based on the distance and thus sacrifices optimality for minimising the overall distance in the tree, the problem tends to be easier to solve for *α→* 0. In fact, in [6], it is shown that finding a linearisation, can be reduced to a maximum weight matching problem, making it polynomial. We can use this situation to find solutions early.

Intuitively, adjacencies that are particularly frequent in a subtree – while not necessarily optimal – are probably not bad candidates as ancestral adjacencies either. We can use these frequencies as weights for Problem 2.4. Concretely, we use the weighted average of the number of occurrences for an adjacency at an internal node as its weight. We then first optimise versions of Problem 2.4 with *α <* 1 before finally optimising with *α* = 1. This ensures on hard instances of the problem that good solutions are found early.

A known heuristic strategy for SPP is to iteratively improve a solution by solving median problems at internal nodes [24]. We adapt this strategy to solving subtrees of the original tree first. In this step, the heuristic solution found in the weighted step is projected onto a subtree, used as an initial solution for the DCJ-SPP restricted to this subtree, and the DCJ-SPP for the subtree is solved. This gives us local solutions, and the lower bound as well as “variable hints” for the ILP solver from this subtree can be carried over to the problem on the full tree.

## 4 Evaluation

We implemented the Swofford-Maddison algorithm to determine marker numbers, the filtering of the solution space and the solving strategies as described in Section 3, integrating the implementation with the ILP developed in [6]. Importantly, we enforce the minimisation of *δ* | *A*^*M*^△ *B*^*M*^| not by including it in the objective function, but by requiring the cost for deleting markers is exactly the minimum determined previously by the Swofford-Maddison algorithm by a series of linear constraints.

We made the software available at https://github.com/gi-bielefeld/spp_dcj_exact. We call this software *SPP-DCJ-exact* in the following.

### 4.1 Simulated Data

We simulated data using Zombi [11]. Since our solution is intended to fit smaller phylogenies with a recent rearrangement history, we used scaled Zombi’s default rearrangement parameters by a scale of 0.2, obtaining rates of 0.2 for duplications, 0.6 for gene losses, and 0.4 for inversions, transpositions and gene originations. By default, we simulated 10 lineages with the root genome containing 100 markers, unless specified otherwise. In the following, we will examine the impact of these parameters on SPP-DCJ-exact.

#### 4.1.1 Performance

##### Comparison with SPP-DCJ

Recall that given the correct marker counts the naive solution to Small Parsimony for Natural Genomes (Problem 2.1) inputs the entire adjacency space into a linearisation problem. The latest ILP solving this problem is *SPP-DCJ-v2*, developed in [6].

Since the space of adjacencies grows primarily with the number of markers, we varied the number of markers at the root genome of the phylogeny for this experiment. To that end, we simulated phylogenies with root genomes of 5 to 30 markers in increments of 5 with 10 samples per step and examined the solving time by gurobi 13.0 set to a time limit of 1 hour. In case no exact solution was found, we instead examined the *relative optimality gap*, that is the ratio (*b− l*)*/b* between the current best solution *b* and the tightest lower bound *l* found by the solver. Recall that what we optimise for is the minimum number of rearrangements in the tree and thus *l* ≥ 0 and a trivial solution is easily found. Therefore, we clamp the optimality gap to be between 0 and 100%. In this experiment, we did not use the solving strategies outlined in Section 3.4, neither for SPP-DCJ-v2 nor for SPP-DCJ-exact. The results are shown in Figure 3.

**Figure 3.**
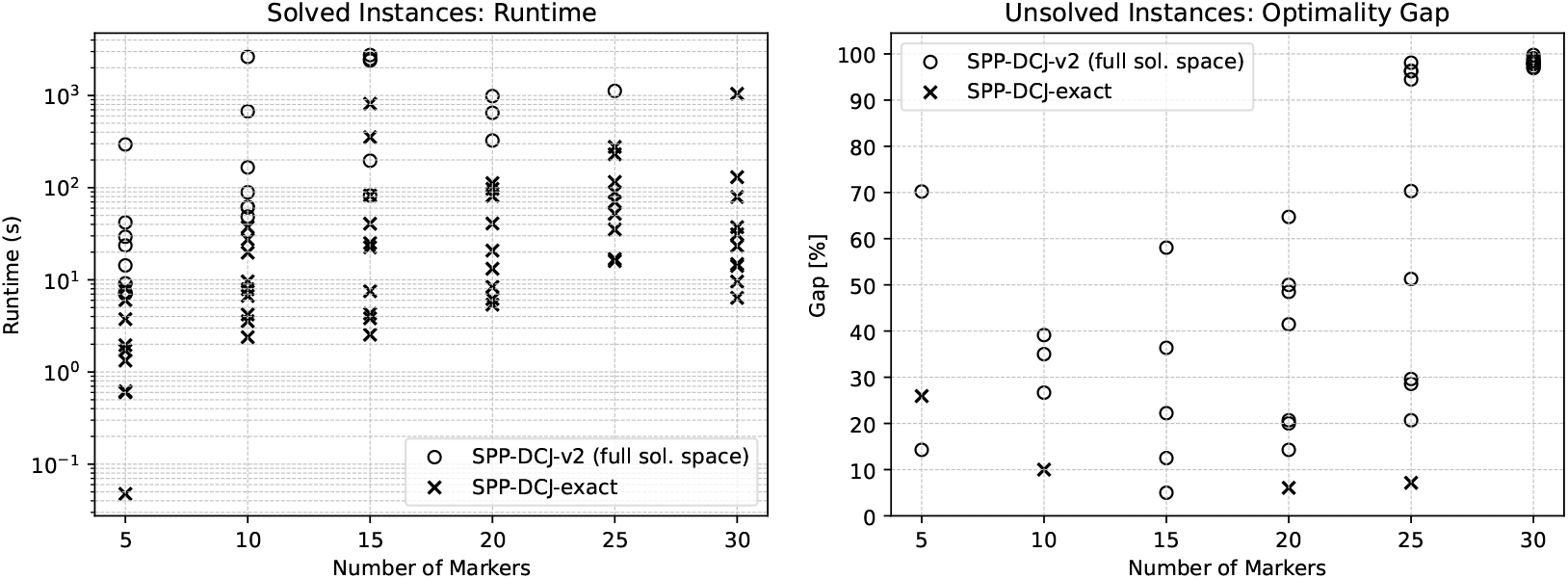
Number of markers vs runtime or optimality gap of SPP-DCJ-v2 and SPP-DCJ-exact as solved by gurobi 13.0 with a time limit of 1 hour (= 3600s). Solved instances are shown on the left, giving their runtime. Instances that exceeded the time limit are shown on the right, instead giving the relative optimality gap reached within 1 hour.

We see that while both ILPs solving times rise with the number of markers, SPP-DCJ-exact outperforms SPP-DCJ-v2, finding exact solutions faster and if no exact solution is found, finding much closer approximations. Notably, almost all SPP-DCJ-v2 instances become unsolvable within the time limit once going above 20 markers, reaching over 90% relative optimality gaps for 30 markers, while the majority of instances of SPP-DCJ-exact remain solvable within 1 hour.

##### Limiting Behaviour

We examined the limits of which problem instances remain solvable using the strategies as detailed in Section 3.4. We evaluated three different settings: *Default*, in which we optimise using only gurobi with a set time limit; *Weights*, in which we allocate 30% of the time limit to solving the problem for *α* = 0.0, 0.1, 0.5 before a final optimisation with the remaining time for *α* = 1; *Weights+Subtrees*, in which we first perform the same weight based solves for 30% of the time limit and then switch to solve subtrees for 30% of the time limit, after which we run the final optimisation.

We performed three experiments, increasing the number of markers, lineages and rearrangements simulated by Zombi. These experiments are discussed in Appendix 1.

To summarise the results, we find a majority of ILPs is solved within an hour up until about 600 markers, 15 lineages and up to 0.3 of the default rearrangement rate in Zombi. We also observe that using the “Weights” or “Weights+Subtree” strategies only starts to positively influence the solution for difficult problem instances and even has an adverse effect for simple instances. Moreover, the “Weights” strategy by itself is typically enough to obtain beneficial effects on optimality and through more time spent on the final optimisation may even outperform the “Weights+Subtrees” strategy.

One possible direction for future research is thus to examine whether these strategies can be improved by using more targeted approach, for example using a specific *α* or solving specific subtrees as well as allocating different ratios of solving time.

#### 4.1.2 Reconstruction Quality

We simulated genomes using Zombi setting rearrangement rates to 0.2, 0.3, 0.4 and 0.5 of the default, generating 20 samples per step and ran SPP-DCJ-exact on a single thread with a time limit of 1 hour on each sample. For this experiment, we used the “Weights” strategy to guarantee a solution. We then examined adjacencies at each internal node separately. Since duplicates complicate the comparison of adjacency sets, we limited ourselves in this analysis to adjacencies involving only non-duplicated markers. On average, each ground truth sample contained 780 such adjacencies.

We then proceeded to evaluate *false positives* (FP), that is such adjacencies that are found in the reconstruction of SPP-DCJ-exact at a given node, but not in the ground truth simulated by Zombi and *false negatives* (FN), such adjacencies that are found in the ground truth, but not in the reconstruction.

The results for all solved instances are shown in Figure 4. We see that the reconstruction for the majority of samples is perfect – the adjacencies reconstructed by SPP-DCJ-exact correspond directly to adjacencies in the ground truth. In fact, all 41 samples without false positives also had no false negatives. Even when there are false reconstructions, the number of incorrectly reconstructed adjacencies observed in this experiment was at most 4 in a single instance.

**Figure 4.**
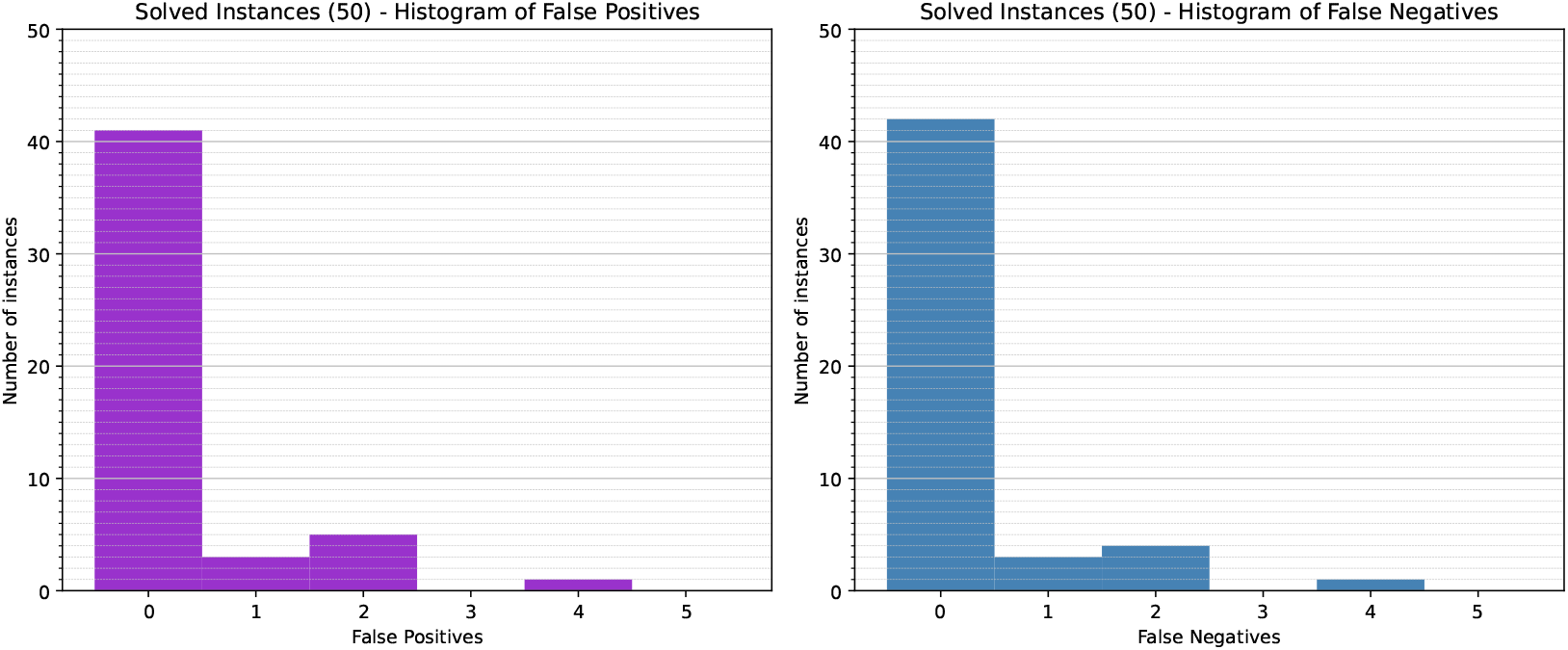
Distribution of errors in reconstructed adjacencies. The histograms show the number of false positives (left) and false negatives (right) on the *x*-axis giving the number of solved instances with a given number of false positives or false negatives on the *y*-axis.

For unsolved instances, we instead examined the relation between the relative optimality gap and the reconstruction quality. To that end, we calculated the *true positives* (TP), i.e. the adjacencies that are found in both the ground truth and the reconstruction at a given node. We then calculated *precision* as *TP/*(*TP* + *FP*), *recall* as *TP/*(*TP* + *FN*) and the *F1-score* as 2*TP/*(2*TP* + *FP* + *FN*). The results are visualised in Figure 5.

**Figure 5.**
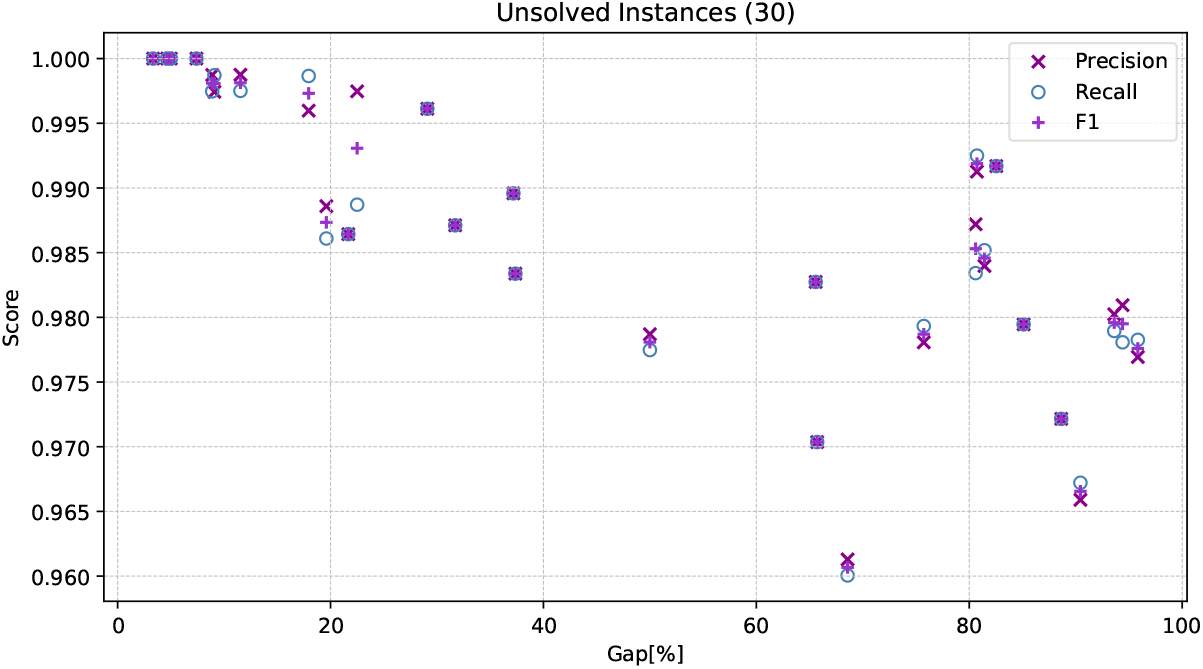
Precision, recall and F1-score in adjacency reconstruction for unsolved instances by relative optimality gap after 1 hour of solving time.

We see that, as expected, solution quality degenerates with a higher gap. However, overall, the quality of solutions remains high with precision and recall for all samples above 0.95, even when approaching a 100% gap. This suggests that SPP-DCJ-exact, at least when the input data has low to moderate numbers of rearrangements, may find optimal or near-optimal solutions, but that the lower bound does not always tighten fast enough to prove optimality.

### 4.2 Real Data

We applied our method to a set of 7 pathogenic plasmids from *Klebsiella pneumoniae*, previously characterised by Biggel *et al* in an epidemiological study of plasmids associated with bovine mastisis [1]. Sampling in this study was constrained to diseased Swiss cows over the course of two years, meaning the plasmid lineages within are recently related and relatively small. We selected the lineage “subcommunity 3” from this study, due to it being the most structurally diverse of the plasmid lineages.

We used MICE [5] to identify synteny blocks to use as markers, using a de Bruijn graph constructed using ggcat [9] with *k*-mer size 31 as input, and filtering out any synteny blocks below 200 bases. Statistics regarding the input genomes generated in this manner are found in Table 1 of Appendix 2. We used the tree topology from Figure 2 in [1] as the input tree. We then applied SPP-DCJ-exact using both weighted and subtree strategies to determine ancestral sequences, using gurobi v11, 8 threads, and setting a time limit of a 12 hours. The best solution encompassed 28 DCJ-indel operations across the tree and was found after 70 minutes. Once the time limit was reached, a lower bound of 21 had been achieved, resulting in a relative optimality gap of 25%. The solution is visualised in Figure 6 with the help of gggenomes package in R [19].

**Figure 6.**
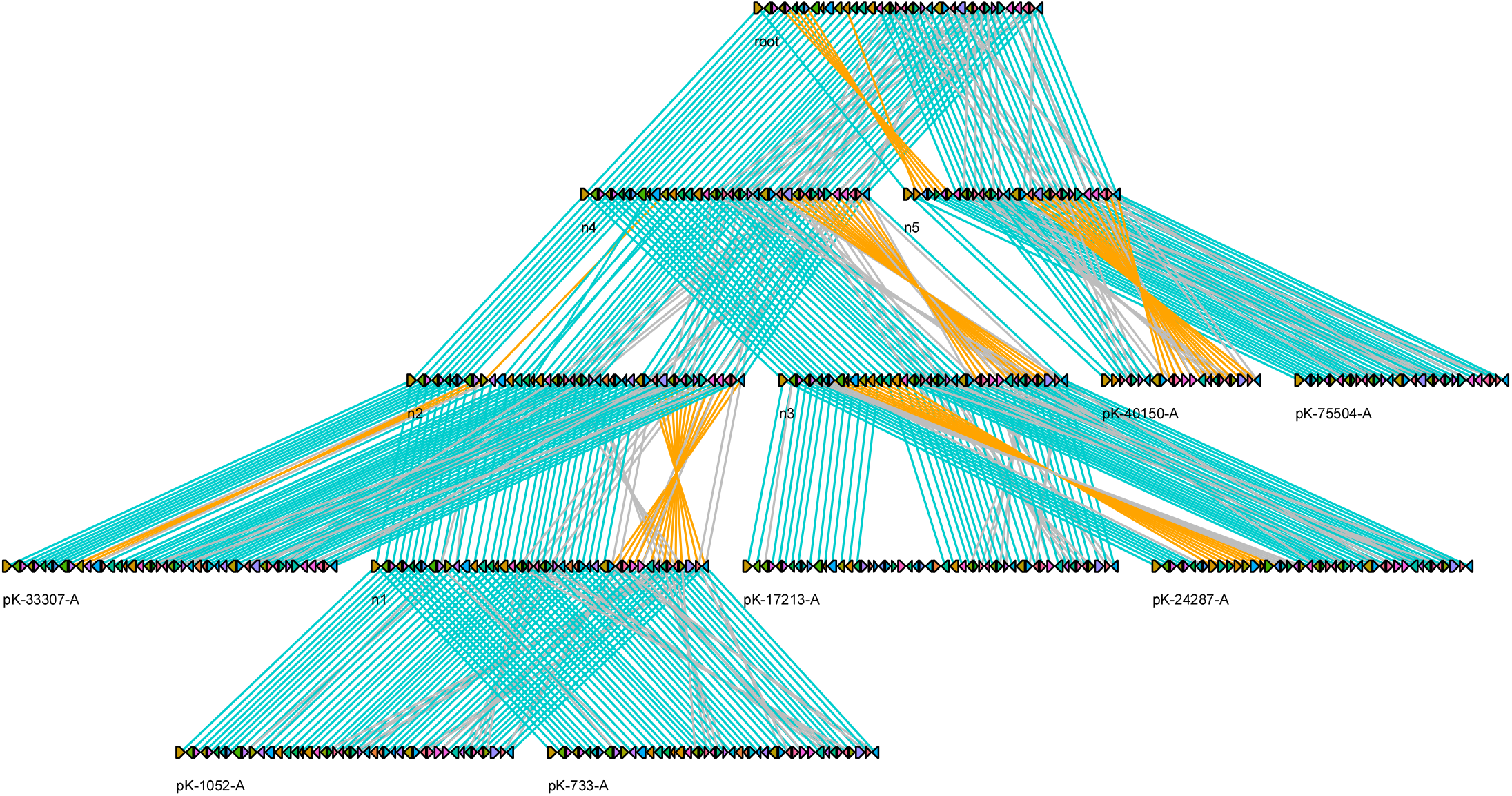
Extant and reconstructed plasmid sequences, arranged according to their tree topology. The same markers across different sequences have the same colour, and marker orientation is shown by arrow direction. Teal links between markers mean they were matched and are oriented the same, orange links between markers mean they were matched and have opposite orientation, while grey lines show links between duplicate markers that were not matched in the final solution. Note that the selection of the root sequence is arbitrary; any intermediary genome between n4 and n5 could be the root, but here we simply choose n4.

All ancestral sequences are circular and unichromosomal, same as the extant plasmids. Their overall size is similar to that of the extant plasmids and, for cherry (n5(pK-40150-A,pK-75504-A)), the parental sequence n5 is larger than both its children. This demonstrates that in practice SPP-DCJ-exact does not “empty out” ancestral genomes, which is a direct result of restoring the triangle inequality. Additionally, there are multiple duplicate markers throughout the tree, including ones nested in rearrangement events spanning multiple markers, but nonetheless the solution parsimoniously matches these markers. This is particularly important for plasmids, as rearrangements are often associated with transposable elements flanked by duplicate genes [20]. From this solution, it is also possible to see that it is the same regions that are undergoing repeated rearrangements, e.g. the markers at the end of all the sequences are inverted multiple times across the whole tree. This is consistent with observations from previous studies of epidemiologically significant plasmids, which usually rely on manual reconstruction of rearrangement events [21, 30]. Overall, we can see in this example that our approach yields biologically plausible ancestral sequences, which can provide insights into the evolutionary history of a plasmid lineage.

## 5 Discussion

We introduced a unified framework for solving the Small Parsimony Problem for natural genomes that requires only extant genomes and the underlying tree as input, while still leveraging the efficient encoding of the solution space using degenerate genomes. In addition, we derived initial theoretical results that substantially restrict the solution space, thereby increasing the number of instances that can be solved exactly or approximated with strong guarantees.

When applied to both simulated data and real data sets with a recent evolutionary history, our approach consistently produces high-quality solutions within reasonable running times, even in cases where optimality was not conclusively proven.

The framework presented here also opens several promising directions for future research. So far, we have investigated only a limited number of simple, though effective, methods to constrain the solution space. A more detailed analysis of the underlying distance measure may allow for an even tighter pruning of degenerate genomes. Moreover, it would be interesting to study scenarios in which differences in marker content are not treated as the dominant term in the distance but can instead be traded off with rearrangement operations (i.e. “*δ <* ∞”). Such an extension could naturally connect our framework to Duplication–Transfer–Loss models of gene tree parsimony, replacing the uniform insertion/deletion cost *δ* with operation-specific costs that distinguish between duplication, transfer, and loss events. Exploring these connections may further broaden the applicability of our approach.

