## Appendix for "Towards a Unified Exact Solution of Rearrangement Small Parsimony for Natural Genomes"

#### 1 Evaluating Limiting Behaviour of Solving Strategies

To assess the performance of the three strategies introduced in Section 4.1.1 (*Default*, *Weights*, and *Weights+Subtrees*), we evaluated their relative optimality gap under a time limit of one hour. We considered three parameters: (i) the number of markers in the root genome, (ii) the number of lineages, and (iii) the number of rearrangements. For each parameter configuration, we generated 10 samples using Zombi. We solved each problem instance with all three strategies, each running on a single thread.

In the first experiment, we increased the number of markers in the root genome from 200 to 1000 in increments of 100. As shown in Figure 1 (A), the optimality gap increases with the number of markers for all three strategies. For small genomes (200–500 markers), most instances are solved optimally within the time limit. Somewhat unexpectedly, the *Default* strategy performed best in this setting, outperforming both *Weights* and *Weights+Subtrees*. A possible explanation is that, for larger genomes, the weighted and subtree-based approaches generate subproblems that themselves become computationally expensive. The additional overhead needed to solve these subproblems may thus "absorb" solving time that would be better used solving the original problem.

In the second experiment, we increased the number of lineages from 11 to 20 in increments of 1. The results are visualised in Figure 1 (B). Across all strategies, the variance in the observed optimality gaps is high. Up to 15 lineages, the majority of instances are solved optimally within the time limit. For 19 and 20 lineages onwards, the *Weights* strategy begins to show an improvement over *Default*. The *Weights+Subtrees* strategy surpasses *Default* only for some instances with a high number of lineages. Notably, however, it exhibits a considerably smaller variance in optimality gaps than both other strategies at these higher values, indicating more stable performance on particularly difficult instances.

In the third experiment, we varied the number of rearrangements. To that end, we scaled duplication rates by $x$, gene loss rates by $3x$, and other rearrangement rates by $2s$, increasing $s$ from 0.2 to 1.0 in increments of 0.1. We call $s$ *rearrangement scale* in the following. As shown in Figure 1 (C), the optimality gap increased sharply with growing rearrangement rates. Most problem instances are solved optimally for $s \leq 0.4$. From $s = 0.7$ onwards, the *Default* strategy almost exclusively exhibits gaps of 100%, indicating that either no solution was found and/or the lower bound could not be tightened further. In the intermediate range $s = 0.6$ to 0.8, *Weights+Subtrees* outperforms *Weights*, but only on some instances. For higher rates of rearrangements, however, the relative optimality gap of all strategies approaches 100%, suggesting the overall time limit of 1 hour is not enough to obtain any significant information about a given problem instance.

Overall, these results indicate that *Weights* and *Weights+Subtrees* are primarily beneficial strategies for solving difficult problem instances. For simple problem instances, the additional preprocessing and decomposition effort appears unnecessary and may even reduce performance. This suggests that further fine-tuning is required, in particular mechanisms for identifying in advance whether solving particular subtrees or applying a weighted optimisation first is likely to improve performance.

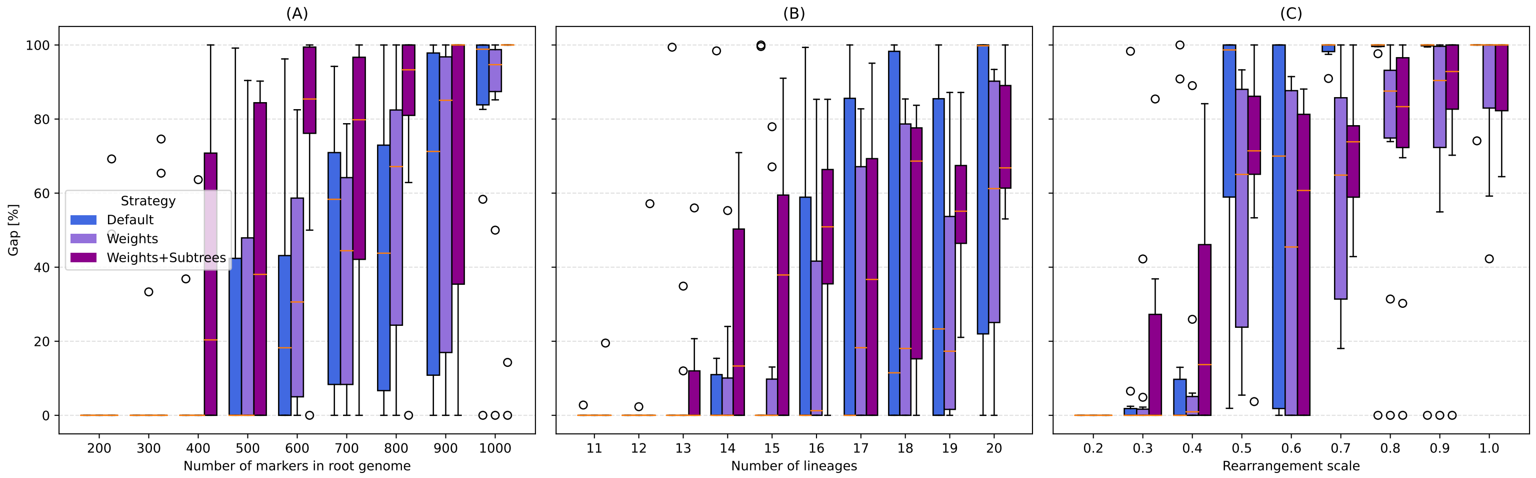


Figure 1: Distribution of optimality gaps for three solving strategies (Default,Weights and Weights+Subtrees) after 1 hour of solving time using gurobi 13.0 in three experiments simulated with Zombi. Relative optimality gaps are visualised as a box plot. (A) The number of markers in the root genome is increased from 200 to 1000 in steps of 100. (B) The number of simulated lineages is increased from 11 to 20. (C) The default rearrangement rate is scaled from 0.2 to 1.0 in increments of 0.1. For each step in each of the experiments 10 samples were generated.

### 2 Plasmid Marker Statistics

| seq_id | length | # singular markers | # duplicate markers |
|---|---|---|---|
| pK-75504-A | 31 | 15 | 8 |
| pK-1052-A | 47 | 29 | 9 |
| pK-33307-A | 47 | 29 | 9 |
| pK-40150-A | 22 | 8 | 7 |
| pK-24287-A | 45 | 19 | 13 |
| pK-733-A | 46 | 28 | 9 |
| pK-17213-A | 53 | 31 | 11 |

Table 1: Summarising statistics of the MICE output used for marker definition. The first column is the plasmid sequence ID, followed by the number of marker occurrences, followed by number of singular markers, and finally the number of duplicate markers.
